# LinCoM: a graph theoretic approach for the production of Linear place fields in Complex Mazes

**DOI:** 10.1101/2020.08.17.253849

**Authors:** Christoforos A. Papasavvas

**Author notes:** Email address (Christoforos A. Papasavvas). Current address: School of Computing, Newcastle University, Newcastle upon Tyne, UK.

## Abstract

Studies on spatial coding and episodic memory typically involve recordings of hippocampal place cell activity while rodents navigate in mazes. Linear place fields serve as reduced representations of the activity of place cells, revealing their spatial preference along the tracks of the maze. Sometimes, the experimental designs include complex mazes with irregular geometries and one or more decision points. Unfortunately, in such complex mazes, the production of linear place fields becomes a non-trivial problem. Here, I present a MATLAB toolbox which implements a graph-theoretic approach for the efficient production of linear place fields in a variety of complex mazes.

## 1. Introduction

The role of hippocampus in navigation and episodic memory is a highly active research area [1]. The indispensable functional element that underlies these cognitive functions is the hippocampal place cell. Place cells fire preferentially at specific locations (i.e., place fields) while an animal moves in space, thus providing spatial coding [2]. Place fields were first reported in an open field area but similar place fields have been recorded on narrow tracks and mazes [3]. Place fields on tracks exhibit directionality; that is, they change depending on the direction of movement along the track [4]. Place cells also encode for the intended destination and learned route when the animal has multiple options in a maze [5, 6].

## 2. Problem and Background

Place cell activity and the video-tracked trajectory of the animal on tracks and mazes are routinely analyzed in tandem for the production of place fields. Subsequent analysis, such as measuring place field size or constructing place cell sequences, relies on the linearization (i.e., transforming the 2D representation to 1D) of the place fields [7, 8]. While linearization is trivial for single linear tracks, it is not trivial in mazes; especially in complex mazes with one or more decision points [9]. A general solution for the production of linear place fields in complex mazes is still missing from the open science toolbox.

The research community would benefit from an algorithm able to efficiently analyze the animal’s trajectory and the place cell activity in order to produce linear representations of each cell’s activity along the different end-to-end paths in a maze (end being a point in a maze where the animal needs to turn back). Notice that the cells’ activity need to be analyzed separately for each end-to-end path (see [5, 6] and illustrative example in Fig. 1). Ideally, the algorithm would be applicable to a variety of mazes made up of interconnected linear tracks. Thus, the problem at hand is, first, to detect individual end-to-end runs (or traversals) in a maze of arbitrary shape, second, to cluster the runs in a path-specific way and, third, to represent the activity of the recorded place cells along the different paths of the maze, that is, to produce a linear place field for each path.

**Figure 1:**
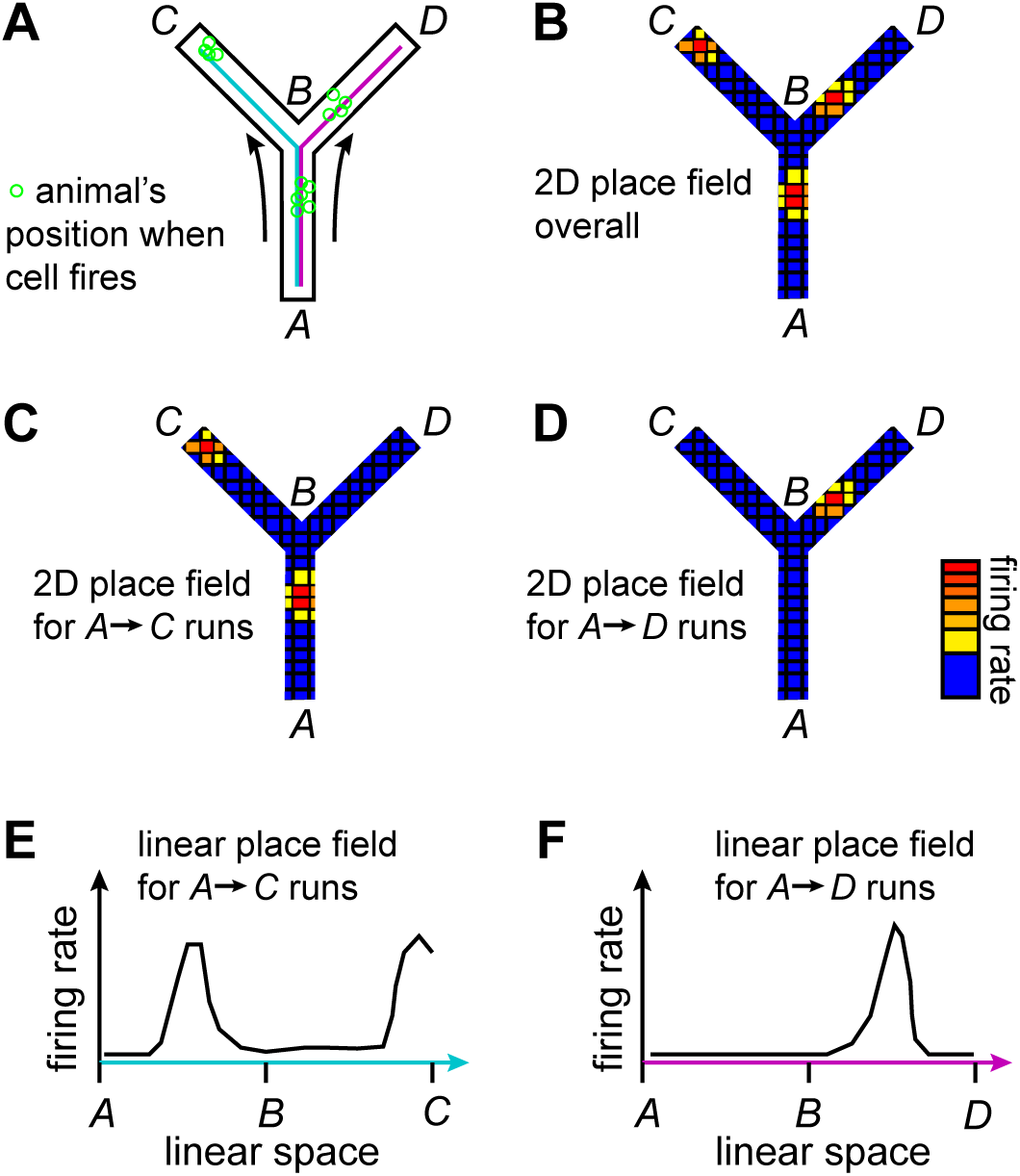
Illustration of the problem and the difference between 2D and 1D (linear) place fields. (A) A rodent moves in a Y maze with one decision point (point B) while a single place cell in hippocampus is being recorded. Starting from point *A*, the animal has the option to follow either the blue path, *A → C*, or the purple path, *A → D*. (B) Overall spiking activity of the cell represented in a 2D firing rate map after multiple *A → C* and *A → D* runs. (C-D) Path specific 2D representations of the activity during *A → C* and *A → D* runs. Notice that the cell fires somewhere between *A* and *B* only when the destination is *C* (indicative of the behavioral goal of the animal [5]). (E-F) After separating the activity between the two paths, linear representations of the cell’s activity are more suitable for subsequent analysis (e.g., construction of place cell sequences [8]).

**Table 1:** Code metadata

| **Nr.** | **Code metadata description** | **Please fill in this column** |
| --- | --- | --- |
| C1 | Current code version | v1.2 |
| C2 | Permanent link to code/repository used of this code version | github.com/cpapasavvas/LinCoM |
| C3 | Legal Code License | MIT |
| C4 | Code versioning system used | git |
| C5 | Software code languages, tools, and services used | MATLAB 2017b |
| C6 | Compilation requirements, operating environments & dependencies | Signal Processing Toolbox, Image Processing Toolbox, Statistics and Machine Learning Toolbox |
| C7 | If available Link to developer documentation/manual | github.com/cpapasavvas/LinCoM/blob/master/README |
| C8 | Support email for questions |  |

Here I present a largely autonomous algorithm and its implementation in MATLAB (LinCoM) for a graph theoretic solution of the problem applicable to a diverse class of user-defined mazes, as opposed to software that are designed for specific mazes (e.g., for W maze, see https://github.com/Eden-Kramer-Lab/MoG_tools). The mazes can have one or more decision points with each one providing two or more alternative paths for the animal to follow. However, the mazes cannot have any cycles; that is, all the branches of the maze must have an end.

## 3. Software Framework

### 3.1. Software Architecture

The software solves the problem by analyzing three types of input data: an image of the maze, the video-tracking data, and the spike-times of one or more cells. The software analyzes the input following the workflow shown in Fig. 2 and outputs the linear place fields for each cell. The image of the maze is only used to create the graph representation of the maze. Using this maze-graph as a reference, the continuous trajectory of the animal (i.e., video-tracking data) is transformed into a discrete trajectory, that is, its trajectory in the maze-graph. Then, the software autonomously detects the runs in the discrete trajectory and clusters them in path-specific clusters. Finally, the spike-times are analyzed in conjunction with the clustered runs in order to produce the linear place fields for each cell.

**Figure 2:**
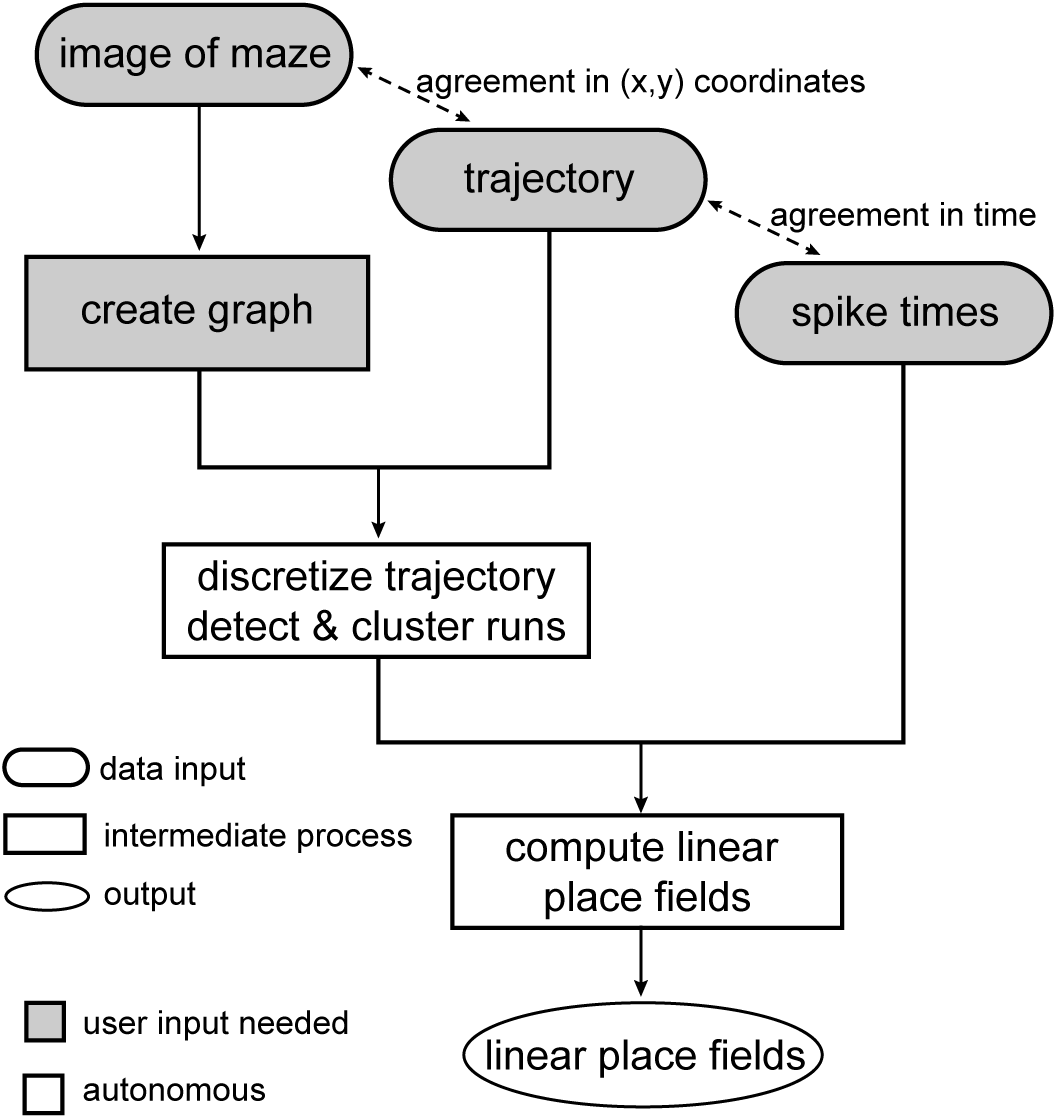
Framework of the software. The framework shows that user involvement is needed only in the initial stages of the whole process while the rest is autonomous. Spatial and temporal relationships between the input elements are indicated with dashed lines. The image of the maze and the trajectory of the animal should refer to the same (*x, y*) plane. Ideally, the image should be taken from the same video that produced the video-tracked trajectory. In addition, the spike-times and the trajectory must be in temporal agreement (i.e., based on the same clock during data collection).

### 3.2. Software Functionality

The software prompts the user to provide all the necessary inputs from the beginning of the process. The software accepts a variety of image and video formats for acquiring an image of the maze. The animal’s trajectory is expected as a *T* × 2 matrix with the (*x, y*) coordinates of the animal for each one of the *T* time-points. The user also needs to provide an *N* × 1 cell array containing the spike-times for each one of the *N* place cells.

The software makes use of interactive features of MATLAB plots to collect some additional user input for the creation of the graph representing the maze. Typical mazes without cycles or open fields, such as T-maze and radial maze, are all supported (see schematics in Supplementary Fig. S1). The software autonomously analyzes the data and outputs a *K × N* cell array containing the *K* place fields of the detected paths for each one of the *N* place cells.

## 4. Implementation

### 4.1. Creation of a spatially embedded graph

The software prompts the user to draw a preliminary graph on top of the image of the maze using interconnected line segments (see example in Supplementary Fig. S2). The line segments are then subdivided into interconnected spatial bins, thus forming a spatially embedded graph *G* (see Fig. 3). The degree of edge subdivision dictates the spatial resolution of the resulting linear place fields and it is set by the user.

**Figure 3:**
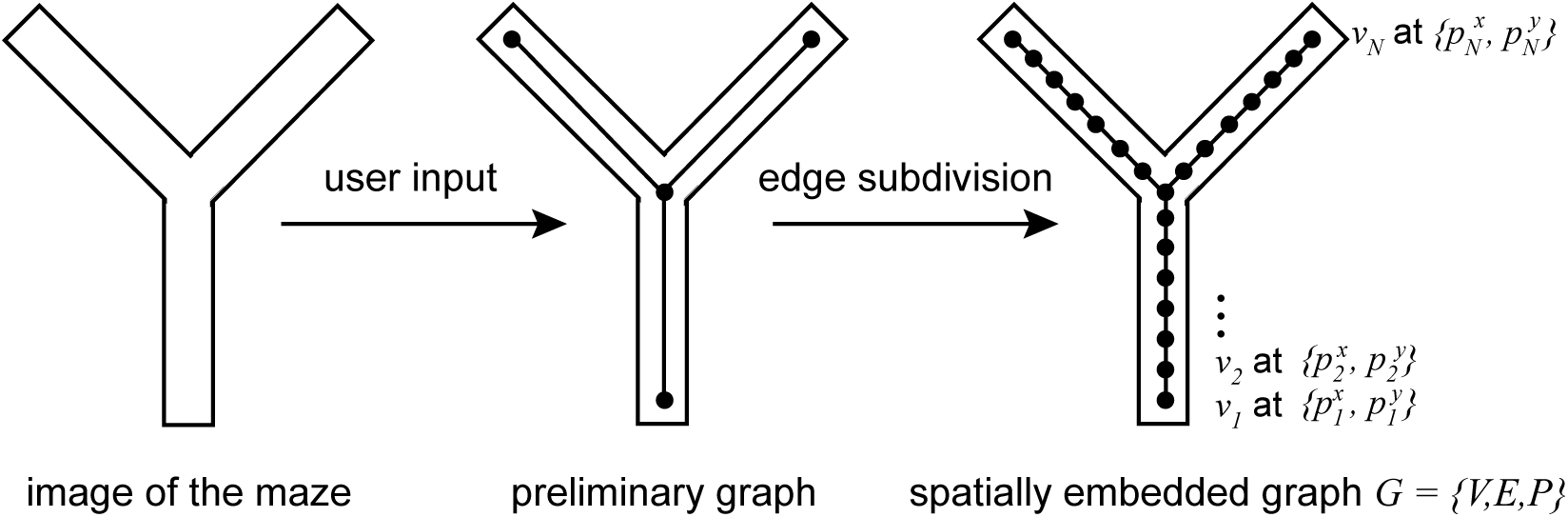
Creation of the spatially embedded graph representing the maze. After a preliminary graph is drawn by the user, the software asks the user for the desired size of the spatial bins and finally creates a spatially embedded graph that represents the maze.

The spatially embedded graph *G* = *{V, E, P}* is defined by an ordered set of *N* nodes *V* = {*{v*_*i*_: *i* = 1 *… N}*}, a set of edges, *E*, and an ordered set of the nodes’ positions in the 2-dimensional space *P* = {*p*_*i*_: *i* = 1… *N*}. Note that *G* is considered to be acyclic and undirected (i.e., a tree in graph theoretic terms), thus the edges have no directionality and there is only one path connecting each pair of nodes. An adjacency matrix *A* is computed from *E*, where the binary value *A*_*ij*_ indicates whether nodes *i* and *j* are connected. The ordering of the nodes in *V* is such that for each node *j* > 2 there is exactly one node *i* < *j* for which *A*_*ij*_ = 1. Given this limitation, the distance matrix *D* is calculated by the dynamic programming algorithm in Supplementary Algorithm S1. The value *D*_*ij*_ indicates the graph-theoretic distance between nodes *i* and *j*. The ordered set of eccentricity values *U* = {*u*_*i*_: *i* = 1… *N*}| is calculated directly from *D* as the maximum values of its rows, that is, the maximum distance of each node from the rest of the graph. The set *Q* of all end-nodes in the graph is also automatically defined.

Graph *G* is supplemented by the user-defined commitment map *C*. The software prompts the user to select interactively a commitment subgraph for each end-node in the maze. During the run detection stage, whenever the trajectory enters a commitment subgraph, the animal is considered to have committed to the corresponding end (see Fig. 4 for an example).

**Figure 4:**
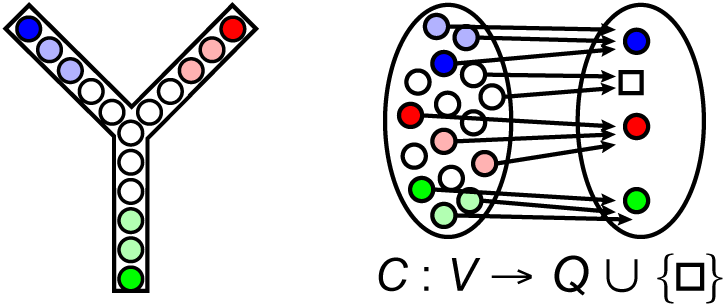
Example of a user-defined commitment map *C*. All nodes in the maze, *v*_*i*_, are mapped either to an end-node, *q*_*i*_, or to a null character.

### 4.2. Discretization of trajectory

In this stage, the continuous trajectory *{X, Y}* is transformed into the discrete trajectory *Z* which is a sequence of graph nodes *z*_1_, *z*_2_,…. For each time-point, the (*x, y*) position of the animal is projected to the nearest node in the graph. The connectivity of the graph is considered during this process such that the projection at time *t*_*i*_ will need to be close, in graph-theoretic terms, to the projection at time *t*_*i*−1_ (see Supplementary Algorithm S3).

### 4.3. Run detection

The algorithm detects individual runs in the maze by using the trajectory eccentricity as a heuristic. It takes advantage of the fact that every time the animal reaches an end in the maze and turns back, the eccentricity of the discrete trajectory *Z* has a local maximum.

First, the discrete trajectory *Z* is reduced in time, resulting in 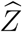, such that there are no consecutive appearances of the same node (i.e., 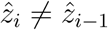). This reduction removes redundant information and simplifies the detection of local maxima. The resulting 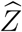 is then expressed in terms of eccentricity producing the trajectory eccentricity *S*. Then, the algorithm finds the local maxima in *S* and saves them in an ordered set *M*, which also stores information about the corresponding node *v* and the corresponding index in 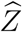. Then the commitment map *C*: *V* → *Q* ∪{□}, where □ is a null character, is applied to *M* so that the nodes that are in a commitment subgraph are mapped to their respective end *q*, while the rest are mapped to □ and removed from the set *M*. Then the algorithm finds all the consecutive pairs of the same end in *M* and, if they are close enough, it removes the one that is less eccentric. Two local maxima are considered to be close enough based on a leeway parameter *L* that allows for a negligible back-stepping of the animal while traveling from one end to another. Finally, the algorithm removes redundant entries in *M* and detects individual runs as subsequences of 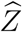 for every consecutive pair of different ends remaining in *M* (see Algorithm 1 and its graphical representation in Supplementary Fig. S3).

### 4.4. Clustering the detected runs

At this stage the algorithm clusters the individual runs in path-specific clusters: one cluster for each unique traverse from one end to another. The user is able to label these clusters with meaningful names relevant to the experimental design (e.g., left-forward, right-backward).

### 4.5. Production of the linear place fields

At the final stage, the software brings together the clusters of detected runs and the spike-times provided by the user. It computes the linear place fields for each cluster and for each place cell. This computation involves the calculation of the time spent (occupation time) and the number of spikes recorded in each spatial bin. The cluster-specific firing rate for each spatial bin is calculated in spikes per second and the linear place field of each cluster is formed by the ordered concatenation of these firing rates (ordered from beginning to end of the corresponding path).

## 5. Illustrative Example

This illustrative example demonstrates the major functions of the software by using a ground truth dataset. This dataset represents a hypothetical scenario where the animal moves in a Y maze (see Fig. 5A). The hypothetical animal executes three times the following sequence of runs: *A* to *C*, *C* to *A*, *A* to *D*, and *D* to *A*. The ground truth data also include spike-times for a single cell with place fields similar to the ones shown in Fig. 1.

**Figure 5:**
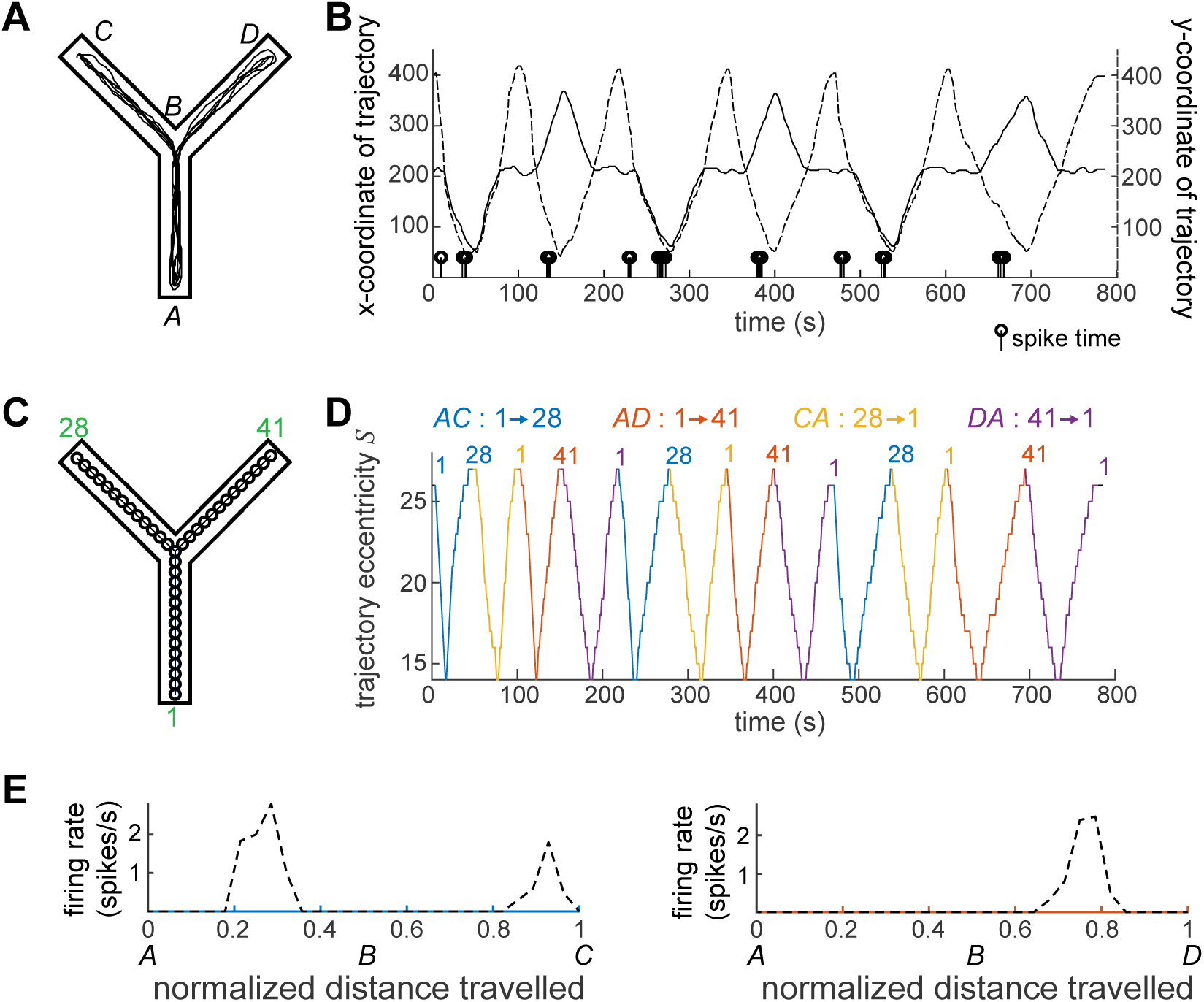
Illustrative example using ground truth data. It demonstrates the most important stages of the process and how the software presents the intermediate and final results. The software: (A-B) accepts a set of raw data; (C) creates and presents the graph; (D) detects, clusters, and presents the runs in time; and finally, (E) produces and presents the place fields for each individual path.

Figure 5A and B provide a visualization of the three input elements: image of the maze, continuous trajectory, and spike-times. After drawing a preliminary graph and setting the size of the spatial bins, the software produced the graph shown in Fig. 5C. Notice that the visualization of the graph includes the node numbers at the three ends. Those numbers appear again in the presentation of the detected runs shown in Fig. 5D. Finally the software produced and presented the linear place fields. Two of them, labeled *AC* and *AD*, are shown in 5E. Notice that the place fields look as expected, since the ground truth dataset was designed to follow the example in Fig. 1.

## 6. Conclusions

LinCoM addresses a problem which increasing number of researchers face in the field of spatial coding in mazes. It provides a general solution for the efficient production of linear place fields in complex mazes with irregular shapes or decision points, thus enabling further quantitative analysis of the place fields and the construction of place cell sequences. Despite the generality of the solution, as it is, the graph-theoretic approach presented here is applicable only to mazes without cycles. The software can potentially be expanded to support mazes with cycles (e.g., 8-figure maze) by using directed graphs as an alternative representation of the maze.

## Supporting information

Supplemental algorithms and figures

## Acknowledgements

I would like to thank the Dragoi lab at Yale School of Medicine for introducing me to this problem. MATLAB R is a registered trademark of The Mathworks, Inc., 3 Apple Hill Drive, Natick, MA 017602098 USA, 508-647-7000, Fax 508-647-7001, www.mathworks.com.

## Algorithm 1 Run Detection

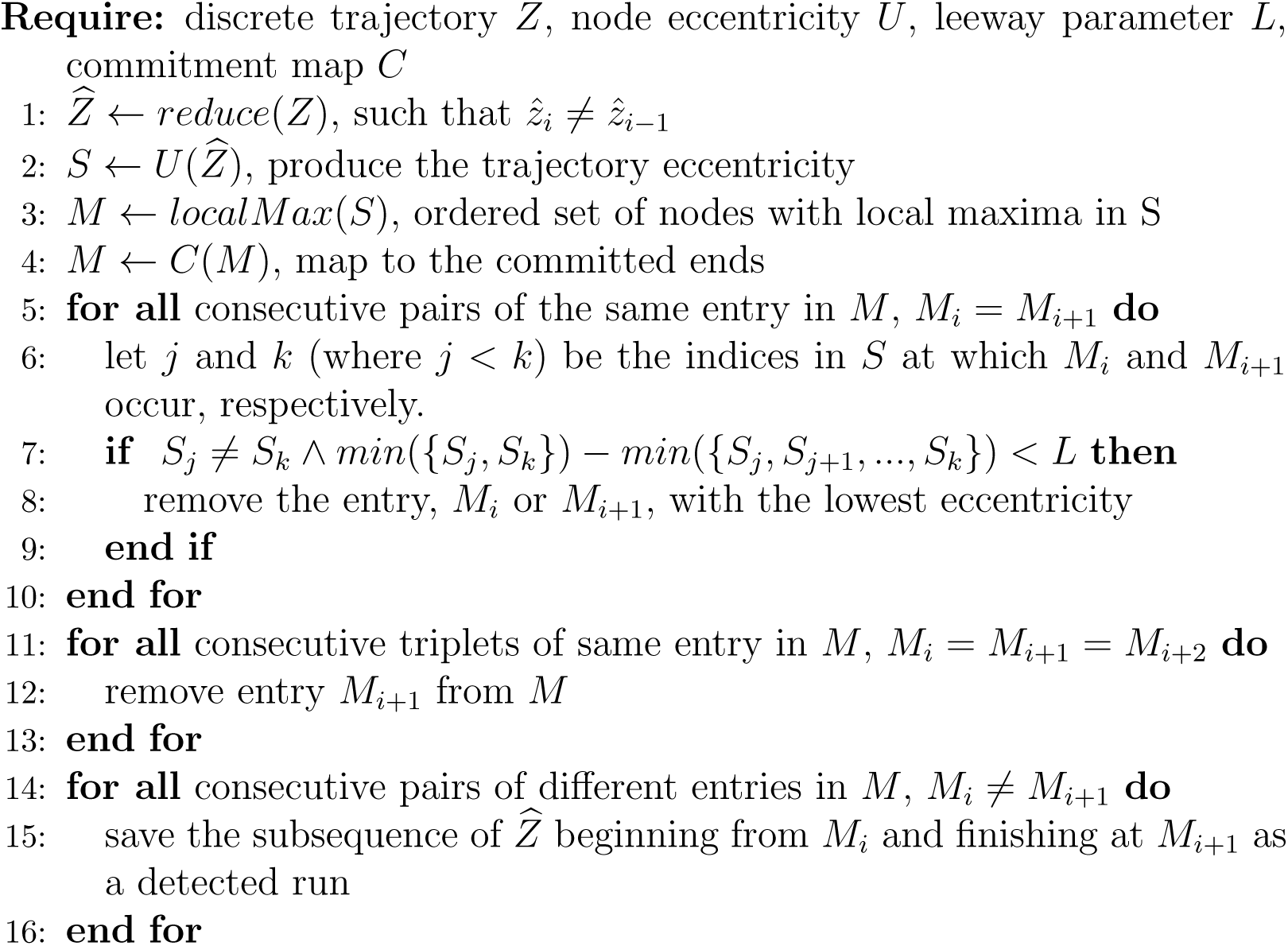

