## Supplemental algorithms and figures for "LinCoM: a graph theoretic approach for the production of Linear place fields in Complex Mazes"

Christoforos A. Papasavvas
Department of Psychiatry, Yale School of Medicine, New Haven, CT, USA

### Supplementary algorithms

**Algorithm S1** Computing Distance Matrix D

**Require:** graph G with adjacency matrix A
**Ensure:** for every $j > 2$ there is exactly one $i < j$ for which $A[i, j] = 1$
**for** j ← 2 to $N$ **do**
    find the only $i < j$ for which $A[i, j] = 1$
    $D[j, 1..j-1] \leftarrow 1 + D[i, 1..j-1]$
    $D[1..j-1, j] \leftarrow D[j, 1..j-1]$
**end for**

**Algorithm S2** find path between start node $v_s$ to finish node $v_f$

**Require:** graph $G = \{V, E, P\}$ with adjacency matrix $A$ and distance matrix $D$
**Ensure:** $v_s, v_f \in V$
path is an ordered set of nodes , initialized with $p = (v_s)$
current node $v_c \leftarrow v_s$
**while** $v_c \neq v_f$ **do**
    find the set J of adjacent nodes to $v_c$, $J = \{v_j : A[v_c, v_j] = 1\}$
    find the node $v_j$ for which $D[v_j, v_f]$ minimizes
    append $v_j$ to path, $p \leftarrow (p, v_j)$
    $v_c \leftarrow v_j$
**end while**

**Algorithm S3** Trajectory discretization

**Require:** continuous cartesian trajectory (X, Y)
**Ensure:** there is a single i for which $i < j$ and $A[i, j] = 1$

compute the euclidean distance of $(x_1, y_1)$ from each node in the graph, $d_k = \|(x_1, y_1) - (p_k^x, p_k^y)\|_2$, $k = 1, .., K$
let $\{v_a, v_b, v_c\}$ be the ordered set of the 3 closest nodes to the initial point $(x_1, y_1)$ with $v_a$ being the closest, based on $d_k$
**if** $A[v_a, v_b] + A[v_a, v_c] + A[v_b, v_c] = 2$ **then**
    $z_1 \leftarrow v_a$
**else**
    ask the user to choose the initial position among $\{v_a, v_b, v_c\}$
**end if**
**for** i $\leftarrow$ 2 to $T$ **do**
    compute the euclidean distance of $(x_i, y_i)$ from each node in the graph, $d_k = \|(x_1, y_1) - (p_k^x, p_k^y)\|_2$, $k = 1, .., K$
    find the closest node to $(x_i, y_i)$, let it be $v_a$
    **if** $D[v_a, z_{i-1}] \leq flex$ **then**
        $z_i \leftarrow v_a$
    **else**
        $z_i \leftarrow z_{i-1}$
    **end if**
    **if** $D[z_i, z_{i-1}] > 1$ **then**
        $(z_{i-1}, z_q, ..., z_i) \leftarrow findPath(z_{i-1}, z_i)$
        $z_i \leftarrow z_q$
    **end if**
**end for**

### Supplementary figures

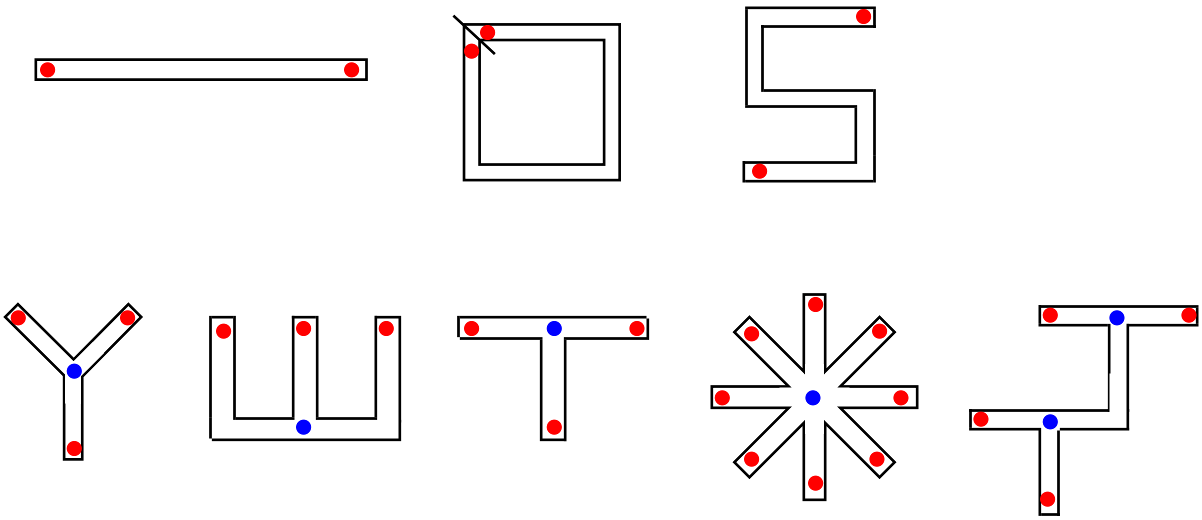


**Figure S1.** Examples of supported mazes. Widely used mazes, such as W maze, radial maze and double T maze, are supported as long as they do not include cycles or open fields (e.i., every branch of the maze needs to have an end, red dots). Alongside the simple mazes at the top, more complex mazes with one or more decision points (blue dots) are also supported.

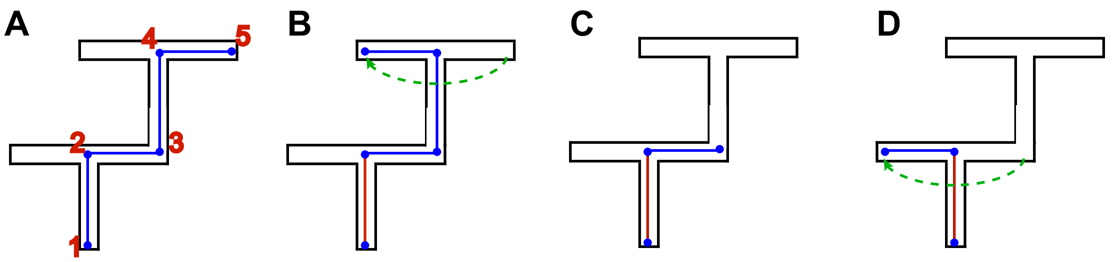


**Figure S2.** Example of drawing interactively a preliminary graph over a double T-maze. (A) Start by tracing the longest end-to-end path. Place successively the five nodes by using left clicks. Right click to finish placing new nodes. (B) Drag and drop node 5 to cover the next end and press enter in MATLAB's command window. (C) Right click on the nodes that are not needed any more to remove them. (D) Drag and drop node 3 to cover the last end and press enter in MATLAB's command window. Note that the first edge drawn (red), between nodes 1 and 2, should always remain stable after the first stage.

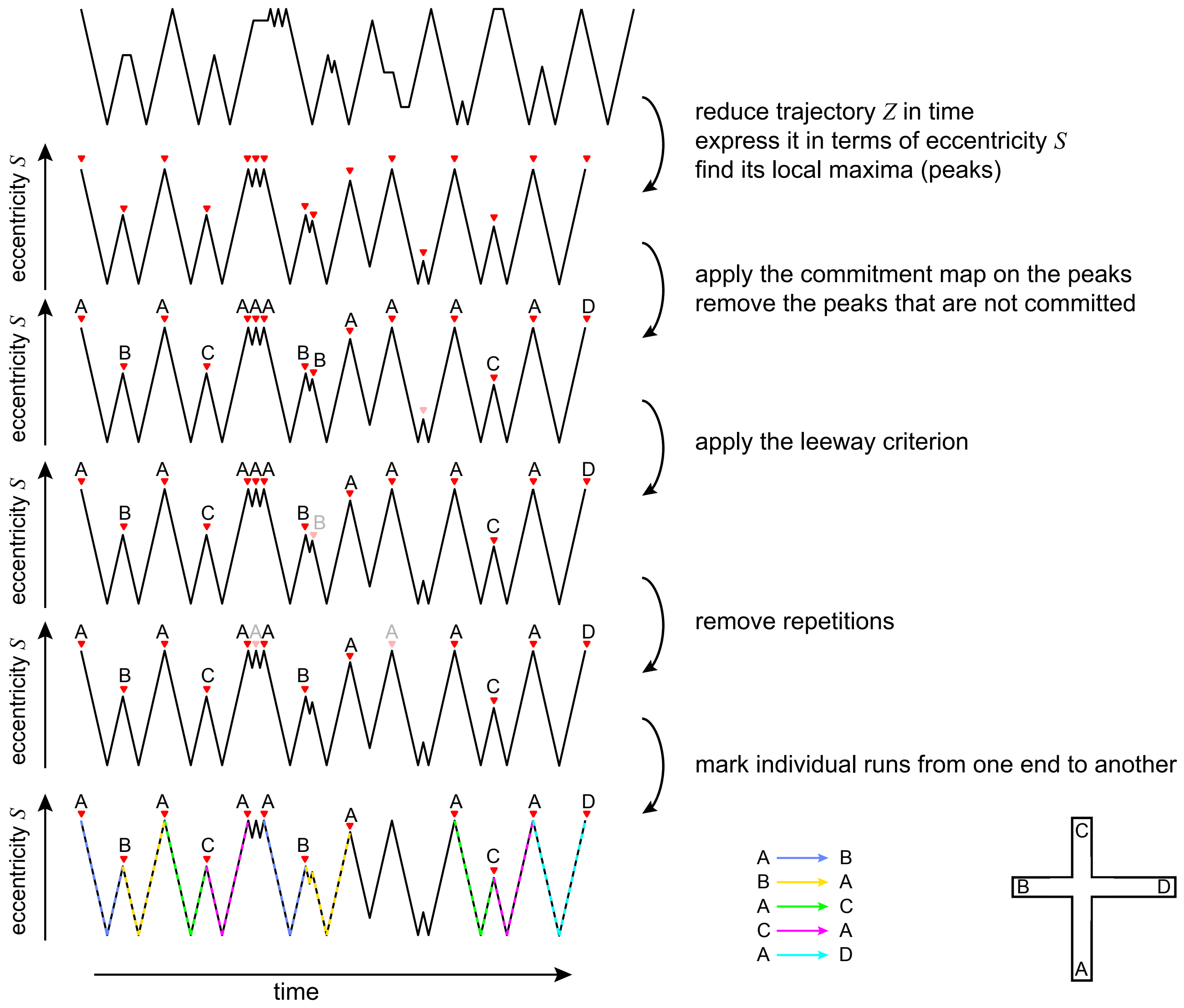


**Figure S3.** Graphical representation of Algorithm 1 for run detection. A 4-arm maze, shown at the lower right corner, is used as an example.
